# Host species background, defence systems, and phage tail gene architecture shape phage infectivity in cystic fibrosis-associated *Achromobacter*

**DOI:** 10.64898/2026.06.29.735440

**Authors:** Anita Tarasenko, Bhavya Papudeshi, Van Nyugen, Susanna R. Grigson, George Bouras, Vijini Mallawaarachchi, Abbey L. K. Hutton, Renee Green, Jack Ramsay, Hamza Hajama, Ana G. Cobián Güemes, Anca M. Segall, Morgyn S. Warner, Sarah K. Giles, Clarice M. Harker, Robert A. Edwards

**Affiliations:** Flinders Accelerator for Microbiome Exploration, College of Science and Engineering, Flinders University, Bedford Park, Adelaide, Australia, 5042; Department of Fundamental Microbiology, University of Lausanne, Lausanne, Switzerland, CH-1005; Microbiology and Infectious Diseases, SA Pathology, Adelaide, South Australia, Australia; DOE Joint Genome Institute, Lawrence Berkeley National Laboratory, Berkeley, CA, USA; Adelaide Medical School, Faculty of Health and Medical Sciences, The University of Adelaide, Adelaide, SA, 5005, Australia; The Department of Surgery - Otolaryngology Head and Neck Surgery, University of Adelaide and the Basil Hetzel Institute for Translational Health Research, Central Adelaide Local Health Network, South Australia, Australia; Department of Biology, San Diego State University, 5500 Campanile Drive, San Diego, CA, 92182, USA; Department of Pathology, University of San Diego, 500 Gilman Drive, MC 0612, La Jolla, San Diego, CA, 92093-0612, USA

**Keywords:** *Achromobacter* spp., Cystic fibrosis (CF), Multidrug resistance (MDR), Phage therapy, Host range

## Abstract

*Achromobacter* species are emerging multidrug-resistant (MDR) pathogens in people with cystic fibrosis. Their increasing resistance has grown an interest in phage therapy as an alternative treatment strategy. However, the factors governing phage susceptibility remain poorly understood, thereby limiting the rational selection of phage candidates. Using 15 strictly lytic *Achromobacter* phages and 7 clinical cystic fibrosis isolates representing *Achromobacter insolitus* and *Achromobacter xylosoxidans,* we demonstrate substantial variation in infection efficiency across all 105 phage-host combinations, variation that could not be discerned from qualitative plaque assays alone. We integrated complete bacterial and phage genomes with quantitative efficiency-of-plating (EOP) assays and lineage-aware Bayesian mixed-effects modelling to show that phage infectivity in *Achromobacter* is governed predominantly by bacterial lineage and strain identity, accounting for 90% of total variance in log-normalised EOP, with individual strains varying substantially in permissiveness irrespective of species membership. After accounting for this lineage structure, no individual defence system, antimicrobial resistance gene class, or phage tail cluster retained a statistically significant independent or interaction association with infectivity. Together, these findings demonstrate that bacterial strain identity is the primary driver of infection outcome. Host defence systems and phage tail-associated genes remain biologically plausible contributors; their independent effect could not be resolved after accounting for lineage structure, indicating that infection outcomes are largely strain-dependent.

This work shifts the question from which individual traits predict infection to how strain lineage and specific host–phage combinations jointly determine infectivity, and argues that quantitative phenotyping of individual phage–host pairs is essential for guiding phage candidate selection and supporting rational cocktail design against multidrug-resistant *Achromobacter* infections in cystic fibrosis.

**Impact statement:** Chronic *Achromobacter* infections in cystic fibrosis are increasingly difficult to treat due to multidrug resistance and biofilm formation. Although phage therapy is a promising alternative, its development is limited by poorly understood and highly variable infectivity. Here, we show that infectivity within a phage host range spans a broad quantitative continuum spanning several orders of magnitude that cannot be captured by qualitative plaque assays. These infection efficiencies are primarily structured by bacterial lineage and strain identity, while the contributions of individual genomic features remain unresolved, given the current sample size. This work provides a framework for predicting phage-host compatibility and supports a shift from empirical screening toward rational, evidence-based phage selection for MDR *Achromobacter* infections.

## 1. Introduction

*Achromobacter* species have emerged as persistent, multidrug-resistant (MDR) pathogens among people with cystic fibrosis (pwCF), and their clinical significance is growing even as highly effective modulator therapies reshape the CF lung microbiome^1–3^. Historically, *Staphylococcus aureus, Pseudomonas aeruginosa*, and *Aspergillus* species dominated chronic CF airway infections. Mucus clearing, facilitated by the widespread adoption of elexacaftor-tezacaftor-ivacaftor and related modulators, has reduced colonisation by these classical pathogens while creating ecological niches for opportunistic species previously considered minor contributors^3^. Among these emerging CF pathogens, *Achromobacter* spp., particularly *Achromobacter xylosoxidans* and *Achromobacter insolitus,* along with *Stenotrophomonas maltophilia*, have been implicated in persistent airway infections associated with accelerated pulmonary decline, treatment failure, and poorer post-transplant outcomes^4,5^. Despite its growing clinical relevance, *Achromobacter* remains among the least mechanistically understood CF pathogens, and treatment options are severely constrained by its intrinsic and acquired resistance across most available antibiotic classes^6^.

The burden of resistance is particularly challenging to manage clinically. Strains routinely carry chromosomally encoded β-lactamases, RND-family efflux pumps, and aminoglycoside-modifying enzymes, and their resistance profiles are further diversified by the horizontal acquisition of mobile resistance genes^4,6^. Clinical isolates frequently display reduced susceptibility to carbapenems, piperacillin-tazobactam, and aminoglycosides, the standard-of-care agents for Gram-negative respiratory infections, leaving few reliable therapeutic options for managing chronic *Achromobacter* colonisation in pwCF. This therapeutic gap has renewed interest in phage therapy as a precision approach for managing MDR *Achromobacter* infections and augmenting conventional antibiotic treatment. Compassionate use cases targeting MDR *Achromobacter* have reported clinical improvement in both CF and post-transplant settings^7,8^, and early-phase feasibility studies of personalised inhaled phage therapy are now underway in CF more broadly, reflecting a shift from anecdotal success toward structured clinical evaluation^9,10^.

Despite this momentum, the promise of phage therapy is constrained by a fundamental mechanistic gap: the driver of phage susceptibility remain poorly understood.

Therapeutic outcomes depend heavily on whether a phage can productively infect and replicate within a target isolate; however, the emergence of resistance and treatment failure remain common challenges. In *Achromobacter,* fewer than fifty phages have been characterised at the genomic level^11–14^, and most studies have assessed infectivity using qualitative plaque assays that classify interactions as permissive or non-permissive^14–16^. While useful for initial screening, these approaches provide limited insight into infection efficiency and cannot readily distinguish highly productive infections from weak or infrequent replication events. As a result, drawing out the genomic features associated with therapeutically relevant infections remains difficult, limiting the rational selection of phages for clinical use.

In other phage–host systems, successful infections are governed by two mechanistic layers. At the bacterial surface, structures such as lipopolysaccharide, capsule, and outer membrane proteins determine adsorption and initial phage recognition, while intracellular defence systems such as restriction–modification, Cyclic-oligonucleotide-based anti-phage signalling systems (CBASS), and abortive infection pathways act as post-entry barriers to block replication^17,18^. Phage host specificity is primarily governed by receptor-binding proteins and tail architecture, with small sequence variations in these regions frequently leading to shifts in host range^19^. However, productive infection depends not only on adsorption but also on successful replication in the presence of intracellular defences, a requirement that must be considered when studying factors that drive infectivity. In *Achromobacter*, it remains unclear which mechanisms determine infection outcome and efficiency. *Achromobacter* species are not uniformly resistant or permissive to phage infection and have highly variable host ranges, even among closely related strains^11,20,21^. However, previous work has relied on small host panels, qualitative plaque assays, or has not sequenced bacterial genomes, making it difficult to determine whether observed differences reflect species-level physiology, strain-specific defence systems, or technical limitations of qualitative readouts^20,22^. The relative contributions of host species identity, individual defence systems, antimicrobial resistance, and phage structural gene architecture to infection efficiency have not been systematically dissected, and whether phage taxonomy or structural gene composition offers more predictive power remains an open question that qualitative host-range data cannot resolve.

Here, we address these gaps using a systematically characterised dataset of *Achromobacter* phage-host interactions comprising complete genomes of 15 strictly lytic *Achromobacter* phages and 7 clinical MDR *Achromobacter* isolates recovered from pwCF. By integrating quantitative efficiency-of-plating (EOP) assays and statistical modelling, we evaluate whether host lineages, defence systems, antimicrobial resistance genes, and phage structural traits independently explain variation in infection efficiency. Together, these findings will support informed phage selection and cocktail design for therapeutic applications.

## 2. Results

### 2.1 Clinical *Achromobacter* isolates represent two species with distinct defence repertoires and AMR profiles

Seven *Achromobacter* isolates were recovered from sputum samples of pwCF receiving care in Adelaide, South Australia, representing chronic airway infections. Whole-genome sequencing resolved the collection into two *Achromobacter* species most commonly associated with cystic fibrosis: *Achromobacter insolitus* (n=4) and *Achromobacter xylosoxidans* (n =3) (Table S1). Phylogenomic reconstruction assigned the clinical isolates into two established species, with neet, aura, cram and vya clustering within *A. insolitus,* and ayb, suz, and jini clustering within *A. xylosoxidans* (Figure 1A). Genome assemblies ranged from 6.3-7.7 Mbp with GC contents of 62.5-68%, consistent with previously reported *Achromobacter* genomes^23–25^. All *A. insolitus* genomes were assembled as complete circular chromosomes using the Hybracter^26^assembly pipeline.

**Figure 1:**
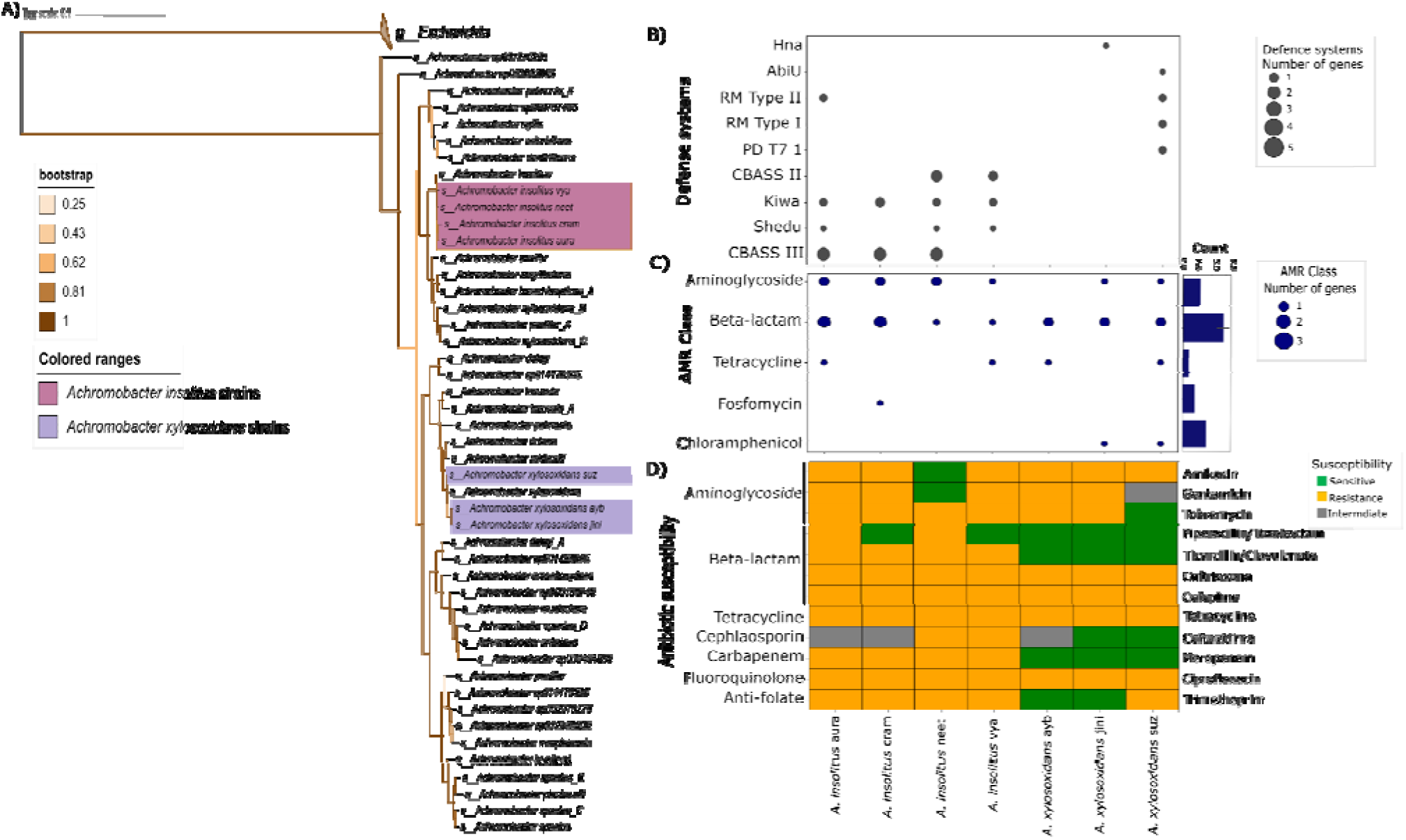
Genomic and antimicrobial resistance profiles of clinical *Achromobacter* isolates. A) Phylogenetic tree reconstruction using GTDB-Tk of *Achromobacter* species, highlighting seven clinical isolates (*A. insolitus* in pink and *A. xylosoxidans* in purple) and 40 reference genomes; *Escherichia* taxa (n=10) used as the outgroup. Branch support is indicated by shading, ranging from lowest (beige) to highest (brown). B) Predicted anti-phage defence systems identified by DefenseFinder and used as putative defence features in downstream analyses. The bubble size represents the number of genes. C) Antimicrobial resistance genes grouped by antibiotic class (AMR class), with the bubble size representing the number of genes. The bar plot to the right represents the average number of unique resistance genes per isolate (presence/absence). D) Phenotypic antibiotic susceptibility profiles for 12 antibiotics (right side of the y-axis), ordered by subclass (left side of the y-axis). Colours indicate sensitivity (green), resistance (orange), and intermediate resistance (grey).

All seven isolates showed MDR phenotypes characteristic of *Achromobacter* recovered from the CF airway, including reduced susceptibility to multiple β-lactams and aminoglycosides (Figure 1D). These phenotypes were supported by genomic profiling, which identified a shared core of resistance genes, including OXA-lineage β-lactamases, RND efflux pumps, and aminoglycoside-modifying enzymes, consistent with previously reported *Achromobacter* resistance signatures (Figure 1C; Tables S2 and S4)^4,6^. Additional strain-specific resistance genes, including *tetC* (aura), *fosA* (cram), and plasmid-borne OXA-5/OXA-10 variants, were also detected, highlighting ongoing diversification of antimicrobial resistance repertoires within the collection.

As antimicrobial resistance profiles alone were unlikely to explain the difference in phage susceptibility, we next examined antiviral defence systems as candidates for genes of infectivity. DefenseFinder^27^ identified 19 defence families across the seven genomes, including CBASS type II and III, Hna, Kiwa, Shedu, multiple restriction-modification systems, and predicted abortive infection pathways (Figure 1B; Table S3). Defence systems varied substantially between strains: *A. xylosoxidans* ayb carried no confidently annotated defence systems, whereas *A. insolitus* neet encoded four, comprising both single-gene systems and multi-component defence pathways (Table S3). Given the known challenges associated with subtype assignment and false-positive predictions across several defence families, these systems were treated as putative genomic features for comparative analyses rather than as experimentally validated antiviral mechanisms^27–30^.

Together, these analyses reveal substantial variation in both antimicrobial resistance and predicted antiviral defence repertoires across clinically relevant *Achromobacter* isolates. This diversity provides a genomic framework for evaluating whether differences in phage infectivity are associated with host-species background, defence-system content, or other bacterial traits.

### 2.2 The *Achromobacter* phage biobank comprises two genomic lineages

To characterise the diversity of *Achromobacter* phages available for therapeutic development, we established a biobank comprising 15 strictly lytic phages, including ten newly isolated phages from South Australian (SA) wastewater and five previously described phages from the San Diego State University (SDSU) Kumeyaay collection (Tables S4 and S5)^11^. All phages produced clear plaques and high-titre lysates, consistent with obligately lytic lifecycles and suitability for therapeutic consideration (Figure 2A).

**Figure 2:**
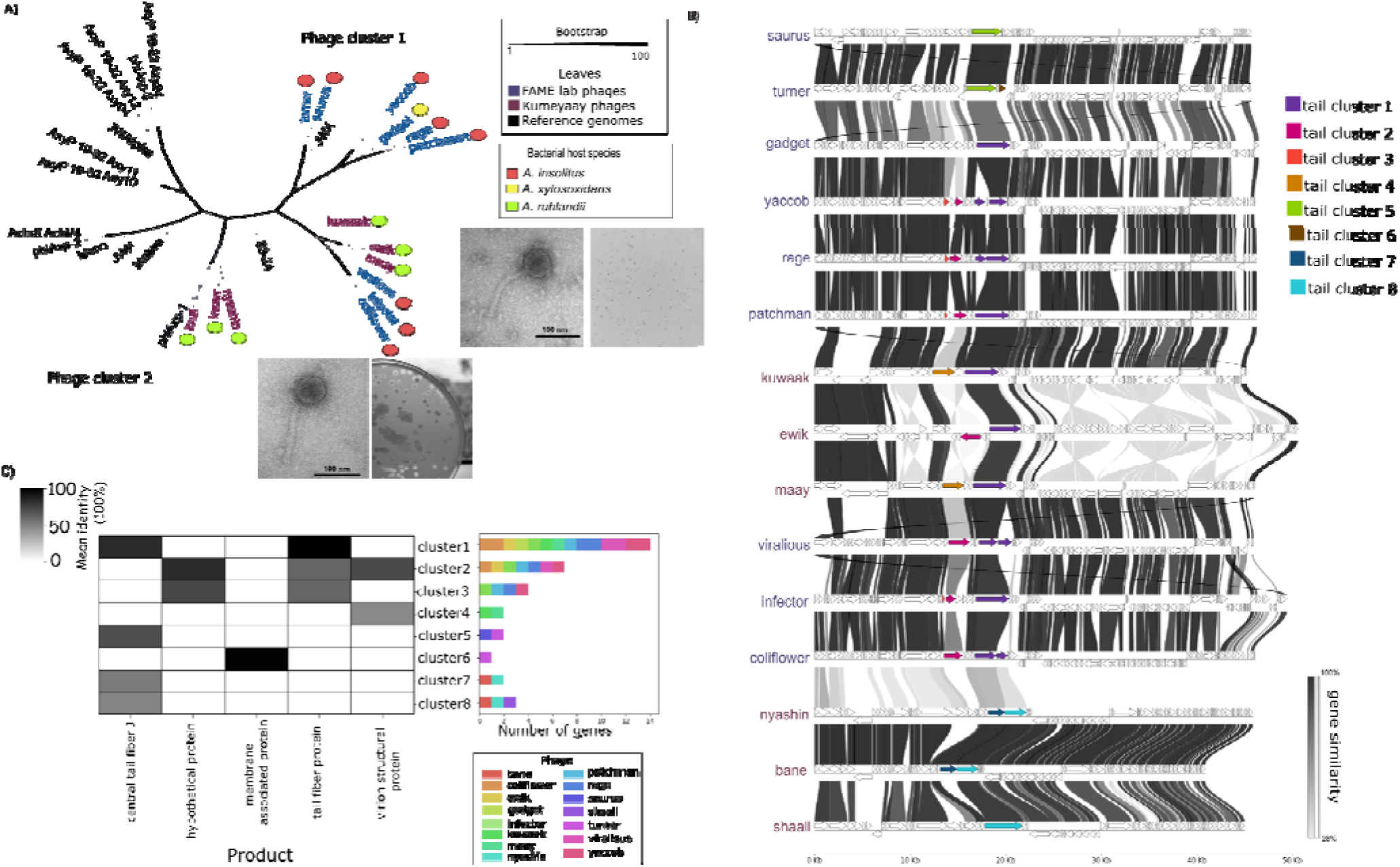
Morphological and genomic characterisation of *Achromobacter* phages. (A) Phage phylogenetic tree inferred from the large terminase subunit (*terL*) gene, with phage highlighted by clusters: Taxa Cluster 1 (*Steinhofvirus*; pink) and Taxa Cluster 2 (unclassified *Siphoviridae*; purple). Coloured circles indicate the bacterial host species on which each phage was isolated: *A. insolitus* (red), *A. xylosoxidans* (yellow), and *A. ruhlandii* (green). TEM micrographs and plaque morphologies of phages coliflower and infector are shown; similar morphologies were observed for all other phages in the collection. (B) Gene-by-gene synteny comparison across the 15 phages. Genes are represented as arrows, and shaded connections indicate amino-acid sequence similarity between homology regions. Tail-associated gene clusters represent the primary regions of genomic variation across the phage collection. C) Heatmap displaying eight tail gene clusters (rows) across five gene product categories (columns): central tail fibre, hypothetical proteins, membrane-associated proteins, tail fibre, and virion structural proteins. Grayscale shading intensity indicates the proportion of genes within each cluster assigned to a given functional category. The barplot panel adjacent to each cluster row displays the total number of genes identified in that cluster for each of the 15 phage isolates.

Transmission electron microscopy (TEM) confirmed siphovirus morphology across all phages, characterised by long flexible tails and isometric capsids (Figure 2A). Although particle dimensions varied modestly among isolates, overall morphology was highly conserved throughout the collection.

Phage genome sizes ranged from 40.8-50.6 kb and encoded 63-107 predicted ORFs with coding densities exceeding 94% (Table S4 and S5). All genomes displayed the canonical modular organisation expected of lytic phages, including DNA replication, morphogenesis, packaging, and lysis. No integrases, recombinases, antimicrobial resistance genes, virulence factors, or putative toxins were detected, supporting their suitability for therapeutic development and consistent with current phage screening frameworks.

Comparative genomics analyses using gene-based phylogenies resolved the biobank into two major genetic clusters (Figure 2A; Figure S1). Cluster 1 comprised ten South Australian phages and three Kumeyaay phages and was most closely related to members of the *Steinhofvirus* genus. Cluster 2 comprised two Kumeyaay phages (Shaaii and Nyashin) together with one South Australian isolate (Bane), forming a distinct, unclassified siphovirus lineage. Notably, phage clustering did not correspond to the host species used for isolation or to geographic origin, indicating no strongly structured genomic relatedness.

Despite overall conservation of genomic architecture, substantial variation was concentrated within the tail-associated regions (Figure 2B). Comparative analysis identified eight tail-gene clusters distributed across otherwise syntenic genomes. These clusters contained predicted receptor-binding proteins, central tail fibre proteins, and other structural components implicated in host recognition and adsorption (Figure 2C). The occurrence of the same functional categories in multiple clusters but with lower sequence identities, central tail fibre J proteins at 48–83% identity and tail fibre proteins at 60–100% identity, further suggests divergence between clusters while retaining functional similarity. The concentration of genomic diversity within these modules suggests that differences in host interaction are more likely to arise from variation in tail-associated genes than from broader differences in phage taxonomy or genome organisation.

Together, these results demonstrate that phage collection comprises two major genomic lineages with broadly conserved genome organisation but substantial diversity in tail-associated genes.

### 2.3 Quantitative EOP assays reveal extensive heterogeneity in *Achromobacter* phage infectivity

We next examined whether this genomic variation translated into differences in infection efficiency using efficiency-of-plating (EOP) assays across clinical *Achromobacter* isolates. This showed substantial variation in *Achromobacter* phage infectivity across the clinical isolate panel (Figure 3). To minimise donor-host-dependent restriction-modification effects, all phage lysates were propagated on the reference strain *A. insolitus* vya prior to testing. Every phage-host combination was assessed in biological triplicate, with low replicate variability observed across several orders of magnitude of infectivity, demonstrating the assay’s reproducibility.

**Figure 3:**
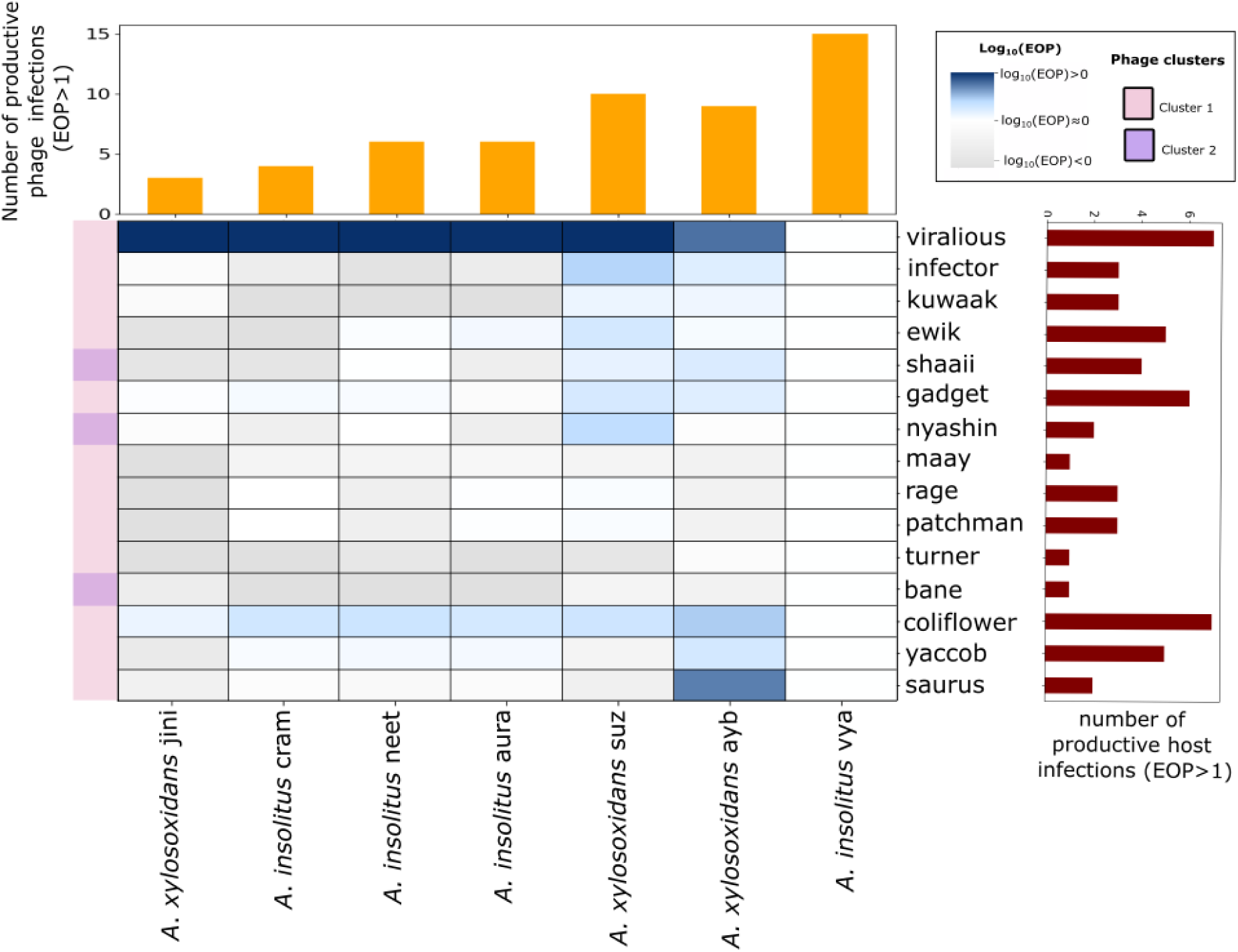
Quantitative efficiency of plating (EOP) profiles of *Achromobacter* phages. Heatmap showing log_10_-transformed EOP values for all phage-host combinations relative to the reference strain *A. insolitus* vya. Positive values indicate higher infectivity than the reference strain; values near zero indicate comparable infectivity; and increasingly negative values indicate reduced infectivity. The upper barplot summarises the number of phages producing productive infections on each bacterial isolate, while the right-hand barplots show the number of hosts productively infected by each phage. The coloured strip adjacent to the heatmap indicates phage genomic cluster assignments. Quantitative EOP measurements reveal substantial heterogeneity in infection efficiency across hosts and phages, spanning several orders of magnitude.

**Figure 4:**
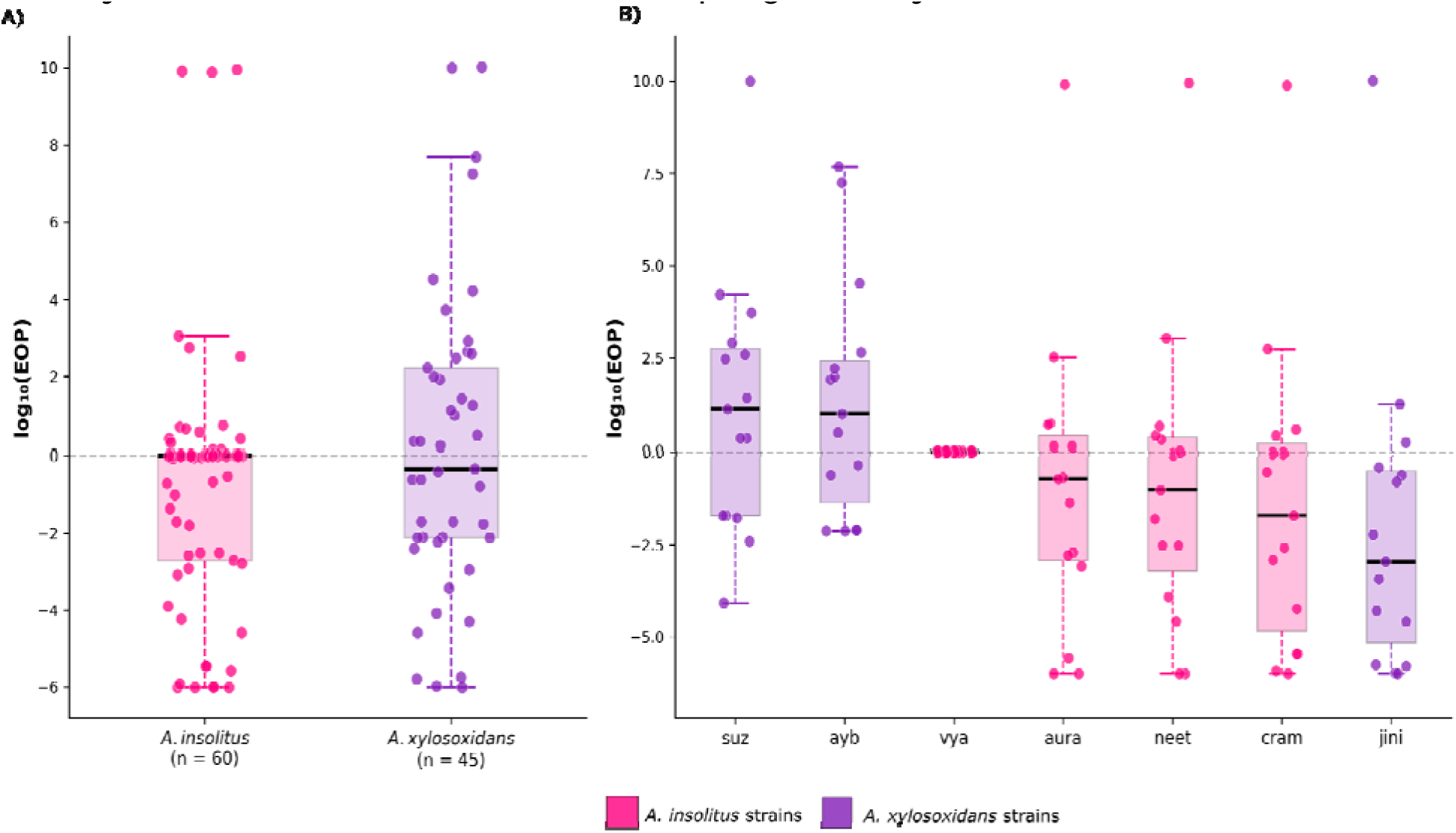
Bacterial strain identity is the dominant driver of phage infection efficiency. (A) Distribution of log_10_(EOP) for all 105 phage-host pairs grouped by host species. The two host species do not differ significantly at the population level (Mann-Whitney U test, p=0.150), reflecting high within-species variation. (B) Distribution of log_10_(EOP) grouped by individual bacterial strain, ordered by median EOP. Each point represents one phage–host pair; boxes show the interquartile range with a median line; the dashed line indicates EOP equivalent to the reference strain (log (EOP) = 0).

EOP provides a quantitative measure of a phage’s ability to adsorb, infect, and complete a productive lytic cycle on a test host relative to a reference strain. Across the *Achromobacter* panel, infectivity spanned several orders of magnitude, demonstrating that phage susceptibility exists along a continuum. Patterns of infectivity varied substantially among hosts. *A. xylosoxidans* ayb and suz supported productive infection by a comparatively large number of phages, whereas *A. xylosoxidans* jini was consistently restrictive and supported productive infection by only a small subset of the collection. Similarly, *A. insolitus* cram displayed markedly reduced susceptibility relative to the other *A. insolitus* isolates (Figure 3). These results indicate that although species-level trends were evident, substantial variation in susceptibility occurred at the level of individual strains.

To compare infection breadth among phages, interactions were classified according to relative infection efficiency. Phages such as coliflower, viralious and gadget exhibited productive infection across the greatest number of hosts, whereas turner, maay, and bane displayed substantially narrower infection profiles (Figure 3). Notably, interactions that produced plaques also exhibited markedly reduced EOP values compared with the reference strain, indicating that plaque formation alone does not necessarily reflect efficient phage replication. These low-efficiency interactions highlight the limitations of binary host-range classification, as infections that appear equivalent in qualitative plaque assays can differ substantially in their capacity to support productive phage amplification^16^.

### 2.4 Lineage-aware statistical modelling best explains the variation in infectivity

To identify bacterial and phage genomic features independently associated with infection efficiency, we fitted a Bayesian linear mixed-effects model with log₁₀(EOP) as the response variable, bacterial defence systems, AMR gene classes, AMR phenotype, and phage tail clusters as fixed effects, and crossed random intercepts for bacterial strain (1|bacteria) and phage identity (1|phage_id). The model converged well across all four chains (maximum R-hat = 1.002) (Figure S2). Variance component analysis revealed that bacterial strain identity accounted for the dominant proportion of variance in log₁₀(EOP) (σ² = 142.53, 90.6% of total variance), with phage identity contributing a smaller but meaningful component (σ² = 9.04, 5.7%), and residual pair-level variation accounting for the remainder (σ² = 5.8, 3.7%). After partitioning this substantial lineage-structured variance into the random effect terms, no individual fixed effect retained a statistically significant independent association with log₁₀(EOP) (all 95% highest density intervals overlapping zero; 0/70 effects significant) (Table S6). Point estimates were in the expected biological directions, like CBASS II (β = −0.19), Hna (β = −0.90), and cluster 6 (β = −6.17), which showed the highest negative associations with EOP, consistent with defence-mediated restriction and cluster-specific infectivity penalties, but credible intervals were wide, spanning up to ±40 log₁₀ units. This reflects insufficient statistical power to resolve individual fixed effects against the dominant background of strain- and lineage-level variance at the current sample size of 7 bacterial strains and 15 phages. Neither main effects nor any of the 48 defence system × tail cluster interaction terms reached significance (Table S6). These results indicate that variation in *Achromobacter* phage infection efficiency is structured primarily by bacterial strain identity, with additional contributions from phage identity.

## 3. Discussion

Phage host range is often treated as a binary phenotype in which a bacterium either supports productive infection or does not. Our analysis of 15 *Achromobacter* phages across 7 clinical CF isolates demonstrates that this framework oversimplifies the complexities of phage-host interactions. Instead, infectivity spans several orders of magnitude and is primarily structured at the level of bacterial species. We show that genomic drivers of infection efficiency appear to be strongly conditioned by host lineage and strain background, although the present dataset lacked sufficient power to resolve the independent effects of individual genomic traits.

Host species identity represented a primary axis of variation in infectivity with *A. insolitus* as broadly higher EOPs than the more restrictive *A. xylosoxidans* (Figure 3). Substantial variation occurred both within and between species, with strain identity explaining approximately 90% of total variance in infectivity. Our finding is consistent with observations across diverse phage-bacteria systems, in which host phylogeny often defines infection boundaries^31,32^. In *Vibrio* and *Staphylococcus*, phage infection has been shown to track genus-level phylogeny, with limited cross-genus infectivity^31,33^. Similarly, in *E. coli,* host phylogeny explained phage infectivity, with additional bacterial and phage traits contributing to finer-scale differences^34^. In *P. syringae*, defence profiles are strongly structured by phylogeny^35^, reinforcing the idea that host evolutionary background shapes susceptibility patterns and must be considered when studying phage-host interactions.

At the same time, host species alone do not fully explain infection outcomes. In *P. aeruginosa*, phage host range is often dominated by variation in bacterial surface receptors rather than by intracellular defence mechanisms, highlighting adsorption-level barriers in some systems^35^. In this collection, broad cross-species infectivity was observed, including phages originally isolated on *A. ruhlandii*^11^ that productively infected both *A. insolitus* and *A. xylosoxidans*. These observations indicate that infection outcomes arise from the interaction between the host’s phylogenetic background and phage-bacterial molecular traits, rather than from host identity alone. On the phage side, tail fibres and receptor-binding proteins play a role in host recognition^19,36,37^. At the structural level, chimeric receptor-binding proteins with sequence-modified RBP modules have been shown to produce phages with altered and predictable host ranges, demonstrating that RBP identity is sufficient to redefine infection specificity independently of overall genome relatedness^38^. Comparison of tail gene clusters across the 15 phages showed substantial sequence diversity despite conservation of key functional roles. Multiple distinct clusters were identified for the same annotated tail genes, indicating that homologous infection-related proteins have diverged considerably at the sequence level. For example, central tail fibre J proteins shared between 48% and 83% sequence identity, while tail fibre proteins exhibited identities ranging from 60% to 100%(Figure 2C). Despite this variation in sequence composition and tail gene architecture, all 15 phages were capable of infecting the same seven bacterial species, but at varying efficiencies (Figure 3). This suggests that considerable sequence divergence can be tolerated within tail-associated proteins while maintaining a conserved host range. This is indicative of the host–phage arms race, in which bacteria evolve resistance to specific phage types and phages counter-evolve to overcome those barriers^39^. These observations suggest that adsorption factors, intracellular defence systems and phage tail architecture may contribute to infection in a lineage-dependent manner. However, their individual effects could not be statistically resolved in the present dataset, highlighting the need for larger collections to disentangle these contributions. From a translational perspective, these findings have direct implications for the design of phage therapy against multidrug-resistant *Achromobacter* infections in cystic fibrosis. Current pipelines rely largely on qualitative host range assays and phage genomic screening, often without incorporating bacterial genomic context. Our results suggest that predictive phage selection will likely require incorporating host lineage and strain background, alongside bacterial and phage genomic features, although larger datasets will be required before these additional predictors can be reliably modelled. As phage biobanks are being established for MDR pathogens, sequencing bacteria and phages, along with quantitative assays, would help build more robust, sensitive frameworks for understanding specific phage-host interactions. In turn, leading to more informed identification of therapeutic candidates and the development of phage cocktails.

Future work should extend this framework to larger and more genetically diverse *Achromobacter* collections, particularly within *A. xylosoxidans*, to refine estimates of within-species variation and improve resolution of the interaction. In addition, because plating efficiency integrates adsorption, replication, and lysis, mechanistic dissection of infection stages using adsorption assays and receptor identification will be necessary to determine which steps are most strongly constrained by specific defence systems.

Extending these analyses into biofilm and *in vivo*-relevant conditions will further establish whether the interaction hierarchy observed here is maintained under clinically relevant environments.

## 4. Conclusions

This study demonstrates that Achromobacter phage–host interactions are structured primarily by bacterial strain identity and broader lineage background. Quantitative infectivity measurements revealed extensive heterogeneity among phage–host combinations that could not be explained by qualitative host-range assays alone.

Although bacterial defence systems and phage structural traits remain plausible contributors to infection outcome, their independent effects could not be resolved after accounting for lineage structure, highlighting the context-dependent nature of Achromobacter phage susceptibility. These findings emphasise the importance of quantitative phenotyping and suggest that larger datasets will be required to disentangle the specific genomic drivers underlying productive infection.

## 5. Methods

### 5.1 Bacterial isolation and culture conditions

Sputum samples from pwCF were collected at SA Pathology (Flinders Medical Centre) under institutional ethics approval and informed consent. Samples were plated on blood agar and incubated at 37°C for 48 h. Morphologically distinct colonies consistent with *Achromobacter* spp. The Flinders Medical Centre Microbiology laboratory, using matrix-assisted laser desorption/ionisation time-of-flight mass spectrometry (MALDI-TOF MS), was streak-purified through three sequential rounds of MacConkey agar followed by Luria-Bertani (LB) agar (1.5% w/v) at 37°C for 16 h. Purified isolates were stored at - 80°C in 20% glycerol. Overnight cultures were routinely grown in LB broth at 37°C with agitation at 180 rpm (Table S1).

### 5.2 Phenotypic antimicrobial susceptibility testing

Phenotypic antimicrobial susceptibility testing was performed using the automated VITEK 2 system (bioMérieux) according to the manufacturer’s instructions. For isolates yielding invalid or indeterminate results on VITEK, susceptibility testing was repeated using gradient diffusion strips (Etest; bioMérieux) or disc diffusion (Oxoid). In these cases, bacterial suspensions were standardised to a 0.5 McFarland inoculum and inoculated onto Mueller-Hinton agar (Edwards Diagnostics). Plates were incubated at 37°C for 18 h prior to interpretation. Results were interpreted using EUCAST clinical breakpoints where available; for agents lacking EUCAST breakpoints, CLSI M100 criteria for non-*Enterobacterales* were applied.

### 5.3 Genomic DNA extraction and sequencing of bacterial isolates

Genomic DNA was extracted from overnight cultures using the NucleoSpin Tissue Mini Kit (Macherey Nagel) according to the manufacturer’s protocol. DNA concentrations and purity were assessed using the Qubit 1x dsDNA HS assay (Invitrogen). Sequencing libraries were prepared from 200 ng of input DNA using the Oxford Nanopore Rapid Barcoding Kit (SQK-RBK114-24). Sequencing chemistry reflected the instruments available at the time of sequencing rather than the experimental design. *A. xylosoxidans* isolates were sequenced on FLO-MIN106 (R9.4.1) flow cells, and *A. insolitus* isolates on FLO-MIN114.24 (R10.4.1) flow cells using a MinION Mk1B device. Raw sequences were deposited in the NCBI Sequence Read Archive (SRA) under the BioProject PRJNA1213176 (Table S1).

### 5.4 Genome assembly, annotation, and resistance/defence profiling

Nanopore FAST5/POD5 files were basecalled using Guppy v6.3.2 in high-accuracy mode, appropriate for each flow cell chemistry (R9.4.1 or R10.4.1)^40^. Adapter sequences were trimmed using Porechop_ABI^41^, and low-quality reads were filtered using fastp^42^. The resulting FASTQ reads were processed within the Baczy v1.0.3^43^ workflow, developed with Snaketool^44^, and executed with Snakemake^45^. Reads were subsampled to approximately 100× coverage using Rasusa v2.2.2^46^. Genome assemblies were generated using Hybracter v0.11.0^26^, which successfully circularised all *A. insolitus* chromosomes. Plasmid reconstruction was performed using Plassembler v1.3.0^47^, and all assemblies were annotated using Bakta v1.10.3^48^.

Antimicrobial resistance genes were identified by integrating predictions from AMRFinderPlus^49^, CARD/RGI^50^ and BV-BRC^51^, with only concordant annotations retained across all three tools (Table S2). Anti-phage defence systems were detected using DefenseFinder v1.2^27^ (Table S3). Because several defence-system families are known to exhibit subtype ambiguity and occasional false-positive predictions, DefenseFinder annotations were treated as putative genomic features for comparative analyses and were not interpreted as experimentally validated defence mechanisms. Fosfomycin resistance was inferred from the presence of a *fos*A-like gene identified during genome annotation; as no EUCAST or CLSI clinical breakpoints are defined for fosfomycin in *Achromobacter,* this annotation was used solely for comparative genomic analyses and not as a clinical resistance classification.

### 5.5 Environmental phage isolation, purification, and propagation

Environmental phages were isolated from wastewater collected at the Glenelg Wastewater Treatment Facility (South Australia) and passed through 0.22 µm filters prior to use. Phage isolation used a soft-agar enrichment approach adapted from Papudeshi *et al.* (2023)^52^. Multiple *Achromobacter* hosts were used for isolation, including *A. xylosoxidans* suz and *A. insolitus* neet and vya strains (Table S4), to maximise recovery of phenotypically and genetically diverse phages. The Kumeyaay phages originally sequenced at SDSU were resequenced at Flinders University in South Australia to verify and refine their genomic data (Table S5).

Briefly, 250 µL of overnight bacterial culture was combined with 500 µL of filtered wastewater, then mixed with 4 mL of molten LB soft agar (0.75% w/v) and poured onto LB agar plates. Plates were incubated overnight at 37°C and inspected for plaque formation. Distinct plaques were re-isolated in three sequential rounds of single-plaque purification to ensure clonal phage populations.

High-titre lysates were prepared by co-incubating purified phages with their respective propagation hosts in LB broth at 37°C with agitation. After overnight incubation, lysates were treated with chloroform, clarified by centrifugation at 4,500 × g for 10 min, and then passed through a 0.22 µm filter. Where required, lysates were concentrated using Vivaspin ultrafiltration units (50 kDa MWCO; Sartorius). Phage titres were determined by standard plaque assays and expressed as plaque-forming units per millilitre (PFU/mL). Lysates were stored at 4°C.

### 5.6 Phage DNA extraction, sequencing, and genome processing

Phage DNA was extracted using the Norgen Phage DNA Isolation Kit with proteinase K digestion and eluted in nuclease-free water. DNA concentrations were quantified using the Qubit 1x dsDNA HS assay (Thermo Fisher Scientific). Sequencing libraries were prepared using the SQK-RBK114-24 rapid barcoding kit (Oxford Nanopore Technologies) and sequenced on Flongle flow cells (R10.4.1). Basecalling was performed using Dorado v0.6.0^53^ with the super-accurate R10.4.1 model, and reads were quality-filtered to retain those with a mean Phred quality score ≥ 10.

Genome assembly and annotation were performed using the Sphae v1.4.6^54^ workflow, which integrates Pharokka^55^ for gene prediction and functional annotation, Phold^56^ for functional annotation using protein structural homology, and Phynteny^57^ for gene synteny-aware functional annotation. Genome circularisation and reorientation were performed using Dnaapler v0.7.0^58^ with genome orientation standardised at the large terminase subunit gene. Functional annotation incorporated PHROG hidden Markov model (HMM) profiles^59^. Gene synteny plots were generated using PyGenomeViz, and orthologous groups were inferred using OrthoFinder v2.5^60^. Particular attention was given to tail-associated structural genes because these regions represented the primary source of genomic variation across the phage collection and were subsequently analysed as candidate drivers of host infectivity. Raw sequencing reads were deposited in the NCBI Sequence Read Archive under the BioProject PRJNA1213176.

### 5.7 Phage comparative genomics and clustering

Pairwise average nucleotide identity (ANI) was calculated using pyANI v0.2.11^61^ under default settings, with ≥95% identity and ≥85% alignment coverage used as preliminary thresholds for cluster delineation. Gene-sharing relationships were assessed using vContact2 v0.11.3^62^ against the RefSeq^63^ viral database. Phylogenetic reconstruction of the large terminase subunit (*terL*) protein was performed by aligning amino acid sequences using MAFFT, followed by maximum-likelihood tree inference with FastTree^64^. The resulting tree was visualised using iTOL^65^

Phage cluster assignments used throughout downstream analyses were based on concordance between ANI relationships, shared gene-content networks inferred by vContact2, and terL phylogenetic reconstruction. Comparative genome visualisation and synteny analyses were used to identify regions of genomic conservation and variation, with particular attention to tail-associated genes, as these regions represented the primary source of genomic diversity across the phage collection.

### 5.8 Host range and efficiency of plating (EOP) assays

Host range profiling and quantitative infectivity measurements were performed using efficiency-of-plating (EOP) assays. Overnight bacterial cultures were mixed with molten LB soft agar (0.75% w/v), poured onto LB agar plates, and allowed to solidify at room temperature. All phage lysates used for host range and EOP analyses were propagated on a single reference strain (*A. insolitus* vya) prior to testing. This strain was selected because it consistently supported robust plaque formation across the phage collection and enabled standardisation of infectivity measurements between hosts while minimising donor-host-dependent restriction-modification effects.

Phage lysates were serially diluted 10-fold in SM buffer from 10 to 10 ¹². For each dilution, 50 µL of phage suspension was mixed with 450 µL of SM buffer and combined with 250 µL of an overnight bacterial culture. Following a 10-minute adsorption period at room temperature, mixtures were added to molten soft agar and overlaid onto LB agar plates. Plates were incubated at 37 °C for 18–24 h, and plaques were enumerated at dilutions yielding countable plaque numbers (typically 30–300 plaques per plate). Phage titres were calculated as plaque-forming units per millilitre (PFU mL ¹) based on the dilution factor and plated volume.

The 10 newly isolated phages and 5 phages from the SDSU Kumeyaay collection^11^ were tested against all 7 clinical *Achromobacter* isolates. Biological triplicates were performed for each phage–host combination^66^. EOP values were calculated as the ratio of phage titre on the test isolate to that on the reference strain (*A. insolitus* vya) and were log10-transformed for downstream analyses. Unless otherwise stated, EOP was treated as a continuous measure of infection efficiency.

Plates were additionally assessed for plaque morphology, clarity, and reproducibility. Only assays producing discrete, countable plaques were included in downstream analyses, and ambiguous outcomes were independently repeated to confirm reproducibility. This quantitative framework enabled discrimination between highly productive infections and low-efficiency interactions that would appear equivalent when assessed solely by plaque presence or absence.

### 5.9 Transmission electron microscopy (TEM)

Phage morphology was examined via negative-stain transmission electron microscopy. Purified lysates were diluted in SM buffer, adsorbed onto plasma-cleaned, carbon/formvar-coated copper grids, stained with 2% uranyl acetate for 30 s, blotted, and air-dried. Imaging was performed on a Tecnai G2 Spirit transmission electron microscope operating at 120 kV. Micrographs were captured with an AMT NanoSprint 15 camera, and particle dimensions were measured using ImageJ v1.53t.

### 5.10 Statistical analysis of genomic traits associated with infection efficiency

To identify bacterial and phage genomic traits associated with infection efficiency, we applied a Bayesian linear mixed-effect model using Bambi v0.15 with PyMC. The model was specified as log₁₀(EOP) ∼ host_species + phage_family + defence subtypes + AMR classes + phenotype_mean + phage tail clusters + subtype: cluster (interaction) + (1 | bacteria) + (1 | phage_id), where host_species and phage_family were included as fixed effects to account for species- and lineage-level baseline differences. Individual defence subtypes and tail clusters were tested as binary fixed effects, and their pairwise interaction terms tested whether the effect of each defence system on EOP depended on the structural gene composition of the infecting phage. Random intercepts were included for bacterial strain (1 | bacteria; n = 7) and phage identity (1 | phage_id; n = 15) as fully crossed random effects, partitioning unexplained variance into strain-level, phage-level, and residual components. Three predictors were excluded prior to fitting due to perfect collinearity: Kiwa (r = 1.00 with host_species), RM Type I and PD-T7-1 (both r = 1.00 with AbiU). Models were sampled using the No-U-Turn Sampler (NUTS) with 4 chains, 2,000 tuning steps, and 2,000 posterior draws per chain (total 8,000 samples), targeting a 0.99 acceptance rate. Convergence was confirmed by R-hat < 1.01 for all parameters. Posterior distributions were summarised as posterior means with 95% highest density intervals (HDI); an effect was considered statistically significant when the 95% HDI excluded zero. Variance components (σ²) were estimated from the posterior means of the random effect standard deviations. All posterior summaries and convergence diagnostics were computed using ArviZ v0.18.

## Acknowledgements

We thank the staff at the Glenelg Wastewater Treatment Facility for their help with environmental sampling. We thank SA Pathology for providing the bacterial isolates and antibiotic sensitivity testing data. Transmission electron microscopy was performed at the Adelaide University Microscopy facility with technical assistance from Cristopher Leigh. Computational resources were provided by Flinders University’s DeepThought cluster^67^, the Pawsey Supercomputing Research Centre and the National Computational Infrastructure (NCI).

## 7. Data availability statement

All raw sequencing data for *Achromobacter* bacterial isolates and their associated phages from this study are deposited in NCBI BioProject under accession PRJNA1213176. Supplementary Table 1 provides BioSamples and SRA accession numbers for bacterial isolates. Supplementary Table 4 lists the SRA and GenBank accession details for the phage isolates. All data will be made publicly available upon publication. During peer review, the data can be accessed using the following reviewer link: https://dataview.ncbi.nlm.nih.gov/object/PRJNA1213176?reviewer=vh51ota528bgvf59p4sjqjr9t0 Code used in this study is available on GitHub: the bacterial assembly and annotation workflow (https://github.com/npbhavya/baczy), the phage assembly and annotation workflow (https://github.com/linsalrob/sphae), and additional analysis scripts (https://github.com/npbhavya/PHI-scripts).

