## Supplementary material for "Host species background, defence systems, and phage tail gene architecture shape phage infectivity in cystic fibrosis-associated *Achromobacter*"

##### Supplementary files:

##### Supplementary Tables S1-S6

##### Supplementary Figures S1-S2

### Supplementary Tables

**Supplementary Table S1:** Sequencing and genome assembly statistics for South Australian *Achromobacter* bacterial isolates

| ***Achromobacter* name** | **jini** | **ayb** | **suz** | **vya** | **neet** | **aura** | **cram** |
| --- | --- | --- | --- | --- | --- | --- | --- |
| **Bioproject ID** | PRJNA1213176 | PRJNA1213176 | PRJNA1213176 | PRJNA1213176 | PRJNA1213176 | PRJNA1213176 | PRJNA1213176 |
| **Biosample ID** | SAMN61177880 | SAMN61177881 | SAMN61177882 | SAMN61177883 | SAMN61177884 | SAMN61177885 | SAMN61177886 |
| **Organism name** | *Achromobacter xylosoxidans* | *Achromobacter xylosoxidans* | *Achromobacter xylosoxidans* | *Achromobacter insolitus* | *Achromobacter insolitus* | *Achromobacter insolitus* | *Achromobacter insolitus* |
| **NCBI ID** | 227019475 | 2211816778 | 2214012089 | 224906884 | 2224603732 #1 | 2224603732 #2 | 2314911183 #3 |
| **Host** | Human | Human | Human | Human | Human | Human | Human |
| **Sample type** | Sputum | Sputum | Sputum | Sputum | Sputum | Sputum | Sputum |
| **Geographical isolation location** | Adelaide, South Australia | Adelaide, South Australia | Adelaide, South Australia | Adelaide, South Australia | Adelaide, South Australia | Adelaide, South Australia | Adelaide, South Australia |
| **Collection date** | 27/08/2022 | 27/08/2022 | 04/04/2023 | `19/09/2023 | 03/09/2022 | 3/09/2022 | 29/05/2023 |
| **Library prep kit** | Rapid sequencing DNA barcoding (SQK-RBK004) | Rapid sequencing DNA barcoding (SQK-RBK004) | Rapid sequencing DNA barcoding (SQK-RBK004) | Rapid barcoding kit 24 V14 (SQK-RBK114-24) | Rapid barcoding kit 24 V14 (SQK-RBK114-24) | Rapid barcoding kit 24 V14 (SQK-RBK114-24) | Rapid barcoding kit 24 V14 (SQK-RBK114-24) |
| **ONT MinION Flowcell** | FLO_MIN106 | FLO_MIN106 | FLO_MIN106 | FLO_MIN114 | FLO_MIN114 | FLO_MIN114 | FLO_MIN114 |
| **Number of reads** | 222,133 | 220,754 | 98,053 | 211,170 | 21,568 | 123,283 | 66,290 |
| **Total length (bp)** | 1,246,472,806 | 1,067,510,810 | 395,743,830 | 1,240,488,647 | 128,835,810 | 768,986,517 | 250,822,579 |
| **Genome coverage** | 197x | 168x | 63x | 180x | 17x | 115x | 37x |
| **Genome coverage (QC bp)** | 0.013 | 0.011 | 0.021 | 0.011 | 0.014 | 0.019 | 0.012 |
| **Total length (assembly bp)** | 6342191 | 6346629 | 6300217 | 6881154 | 7710756 | 6697348 | 6720574 |
| **Chromosome length (bp)** | 6342191 | 6346629 | 6300217 | 6762963 | 7270482 | 6722954 | 6722594 |
| **Plasmid length** | 0 | 0 | 0 | 0 | 2 | 1 | 2 |
| **Number of contigs** | 1 | 1 | 1 | 2 | 9 | 2 | 6 |
| **GC content (%)** | 67.8% | 67.7% | 67.8% | 64.7% | 62.5% | 65% | 65% |
| **tRNA counts** | 58 | 60 | 69 | 61 | 86 | 61 | 60 |
| **tmRNA counts** | 0 | 1 | 1 | 2 | 2 | 1 | 2 |
| **rRNA counts** | 10 | 10 | 10 | 13 | 16 | 13 | 13 |
| **ncRNA counts** | 8 | 7 | 7 | 9 | 11 | 7 | 7 |
| **nc regions** | 12 | 11 | 12 | 17 | 19 | 19 | 18 |
| **CRISPR arrays** | 0 | 0 | 0 | 0 | 0 | 0 | 0 |
| **CDS** | 7,327 | 7,490 | 6,009 | 8,284 | 7,905 | 8,454 | 8,687 |
| **Pseudogenes** | 0 | 0 | 0 | 0 | 0 | 0 | 0 |
| **Hypothetical proteins** | 3,988 | 4,262 | 1,308 | 4,763 | 4,060 | 5,113 | 5,594 |
| **Signal peptides** | 0 | 0 | 0 | 0 | 0 | 0 | 0 |
| **AMRFinders** | 4 | 3 | 5 | 12 | 12 | 11 | 9 |
| **Defense systems** | 1 | 0 | 4 | 3 | 4 | 4 | 3 |
| **Prophages** | 3 | 2 | 3 | 4 | 3 | 3 | 0 |

##### **Supplementary Table S2:** Summary of antimicrobial resistance (AMR) genes detected in *Achromobacter* isolates using NCBI AMRFinder

| **Isolate** | **Species** | **AMR genes detected** | **Class** | **Subclass** | **Plasmid Chromosome** | **notes** |
| --- | --- | --- | --- | --- | --- | --- |
| jini | *A. xylosoxidans* | *catB*, *aph*(3'), *ampC*, *blaOXA* | Phenicol, Aminoglycoside, Beta-lactam | cat, APH, C, OXA | Chromosome | blaOXA-114 partial hit |
| ayb | *A. xylosoxidans* | *ampC*, *blaOXA* | beta-lactam | C, OXA | Chromosome | blaOXA (OXA-114 family) |
| suz | *A. xylosoxidans* | *catB*, *aph*(3'), *ampC*, *blaOXA* | Phenicol, Aminoglycoside, Beta-lactam | cat, APH, C, OXA | Chromosome | blaOXA (OXA-114 family) |
| vya | *A. insolitus* | *ampC*, *aph*(3')-II | Beta-lactam, Aminoglycoside | C, APH-II | Chromosome | Both chromosomal hits |
| neet | *A. insolitus* | *ampC*, *aph*(3')-II, *aac*(6') | Beta-lactam, Aminoglycoside | C, APH-II, AAC | Chromosome | High coverage hits |
| aura | *A. insolitus* | *ampC*, *aph*(3'), *aac*(6'), *blaOXA* (x2) | Beta-lactam, Aminoglycoside | C, OXA, APH | Chromosome + plasmid | blaOXA detected on plasmid |
| cram | *A. insolitus* | *fos*, *aph*(3'), *aac*(6'), *blaOXA* (x3) | Fosfomycin, Beta-lactam, Aminoglycoside | fos, OXA | Chromosome + plasmid | Multiple blaOXA variants |

##### **Supplementary Table S3:** Predicted bacterial defence systems in *Achromobacter* isolates

| **Isolate** | **Species** | **Defence system detected** | **Subtypes** | **No. of genes** | **Notes** |
| --- | --- | --- | --- | --- | --- |
| jini | *A. xylosoxidans* | Hna | Hna | 1 | Single Hna gene |
| ayb | *A. xylosoxidans* | - | - | 0 | No system detected |
| suz | *A. xylosoxidans* | AbiU, RM, PD-T7-1 | RM-Type I, RM-Type II | 7 | Multiple RM and AbiU present |
| vya | *A. insolitus* | CBASS, Kiwa, Shedu | CBASS-II | 6 | Three distinct systems |
| neet | *A. insolitus* | CBASS, Kiwa, Shedu | CBASS-II, CBASS-III | 12 | Two CBASS subtypes present |
| aura | *A. insolitus* | CBASS, Kiwa, Shedu | CBASS-II, RM-Type II | 10 | Combination of RM + CBASS |
| cram | *A. insolitus* | CBASS, Kiwa, Shedu | CBASS-III | 9 | Three systems detected |

##### **Supplementary Table S4:** Metadata and genomic characteristics of *Achromobacter* phages isolated from Adelaide wastewater in September 2023

| ***Achromobacter* name** | **Rage** | **Yaccob** | **Patchman** | **Viralious** | **Turner** | **Saurus** | **Coliflower** | **Infector** | **Gadget** | **Bane** |
| --- | --- | --- | --- | --- | --- | --- | --- | --- | --- | --- |
| **Biosample IDs** | SAMN463214773 | SAMN46321774 | SAMN46321775 | SAMN46321776 | SAMN46321777 | SAMN46321778 | SAMN46321779 | SAMN46321780 | SAMN46321781 | SAMN60942879 |
| **SRA ID** | [SRR36349813](https://dataview.ncbi.nlm.nih.gov/object/SRR36349813) | [SRR36349812](https://dataview.ncbi.nlm.nih.gov/object/SRR36349812) | [SRR36349811](https://dataview.ncbi.nlm.nih.gov/object/SRR36349811) | [SRR36349810](https://dataview.ncbi.nlm.nih.gov/object/SRR36349810) | [SRR36349809](https://dataview.ncbi.nlm.nih.gov/object/SRR36349809) | [SRR36349808](https://dataview.ncbi.nlm.nih.gov/object/SRR36349808) | [SRR36349807](https://dataview.ncbi.nlm.nih.gov/object/SRR36349807) | [SRR36349806](https://dataview.ncbi.nlm.nih.gov/object/SRR36349806) | [SRR36349805](https://dataview.ncbi.nlm.nih.gov/object/SRR36349805) | SRR39271494 |
| **Organisms name** | *Achromobacter* phage Rage | *Achromobacter* phage Yaccob | *Achromobacter* phage Patchman | *Achromobacter* phage Viralious | *Achromobacter* phage Turner | *Achromobacter* phage Saurus | *Achromobacter* phage Coliflower | *Achromobacter* phage Infector | *Achromobacter* phage Gadget | *Achromobacter* phage Bane |
| **Host** | *A. insolitus* neet | *A. insolitus* neet | *A. insolitus* neet | *A. insolitus* neet | *A. insolitus* vya | *A. insolitus* vya | *A. insolitus* neet | *A. insolitus* neet | *A. xylosoxidans* suz | *A. insolitus* neet |
| **Library prep kit** | Rapid barcoding kit 24 V14 (SQK-RBK114-24) | Rapid barcoding kit 24 V14 (SQK-RBK114-24) | Rapid barcoding kit 24 V14 (SQK-RBK114-24) | Rapid barcoding kit 24 V14 (SQK-RBK114-24) | Rapid barcoding kit 24 V14 (SQK-RBK114-24) | Rapid barcoding kit 24 V14 (SQK-RBK114-24) | Rapid barcoding kit 24 V14 (SQK-RBK114-24) | Rapid barcoding kit 24 V14 (SQK-RBK114-24) | Rapid barcoding kit 24 V14 (SQK-RBK114-24) | Rapid barcoding kit 24 V14 (SQK-RBK114-24) |
| **Number of reads (RAW)** | 23,002 | 28,440 | 23,932 | 21,451 | 9,661 | 14,622 | 43,424 | 8,415 | 25,354 | 701 |
| **Total length (RAW bp)** | 23,603,004 | 68,752,675 | 56,315,363 | 46,532,516 | 20,041,550 | 27,009,549 | 210,426,113 | 40,506,449 | 183,050,909 | 3,752,012 |
| **Total length (assembly bp)** | 46,104 | 46,120 | 46,120 | 46,129 | 45,697 | 45,704 | 46,110 | 49,417 | 46,469 | 40,845 |
| **GC content (%)** | 56% | 56% | 56% | 56% | 56% | 56% | 56% | 56% | 56% | 56% |
| **CDS** | 89 | 89 | 86 | 88 | 91 | 86 | 93 | 107 | 91 | 81 |
| **Hypothetical proteins** | 50 | 48 | 47 | 51 | 57 | 53 | 54 | 69 | 59 | 58 |
| **Closest reference genome** | Achromobacter phage vB_AxyS_19-32_Axy16 | Achromobacter phage vB_AxyS_19-32_Axy16 | Achromobacter phage vB_AxyS_19-32_Axy16 | Achromobacter phage vB_AxyS_19-32_Axy16 | Achromobacter phage JWX | Achromobacter phage JWX | Achromobacter phage vB_AxyS_19-32_Axy16 | Achromobacter phage AMA1 | Achromobacter phage AMA1 | Achromobacter phage vB_AxyS_19-32_Axy20 |

**Footnote:** All phages produced circular, clear plaques with no halo formation, though plaque size varied among isolates. Plaque diameters were measured for a subset of phages during TEM preparation: Turner (5–7 mm), Coliflower (1–2 mm), Infector (3–4 mm), and Gadget (5–6 mm). The subset of phages visualised using TEM was classified within the *Siphoviridae* family and exhibited isometric heads. Measured head dimensions and tail lengths were as follows: Turner (head length/width: 66 nm; tail length: 151 nm), Coliflower (head length/width: 56.8 nm; tail length: 141 nm), Infector (head length/width: 62.6 nm; tail length: 140 nm), and Gadget (head length/width: 713 nm; tail length: 158 nm).

##### **Supplementary Table S5:** SDSU *Achromobacter* phage morphology and characteristics from [Cobián Güemes et al., 2023](https://doi.org/10.3390/v15081665)

| ***Achromobacter* name** | **Nyashin** | **Kuwaak** | **Shaaii** | **Maay** | **Ewik** |
| --- | --- | --- | --- | --- | --- |
| **Bioproject ID** | PRJNA123176 | PRJNA123176 | PRJNA123176 | PRJNA123176 | PRJNA123176 |
| **BioSample ID** | SAMN6321782 | SAMN46321783 | SAMN60942874 | SAMN46321785 | SAMN46321787 |
| **SRAs** | SRR36349722 | SRR36349721 | SRR39271495 | SRR36349719 | SRR36349717 |
| **Organisms name** | *Achromobacter* phage Nyashin | *Achromobacter* phage Kuwaak | *Achromobacter* phage Tuull | *Achromobacter* phage Maay | *Achromobacter* phage Ewik |
| **Host** | *Achromobacter ruhlandii* | *Achromobacter ruhlandii* | *Achromobacter ruhlandii* | *Achromobacter ruhlandii* | *Achromobacter ruhlandii* |
| **Isolation source** | Influent water, Cardiff, CA | SDSU fishpond | Influent water, Cardiff, CA | Quality Lab Escondido, Ca | Lake Murray |
| **Geographical isolation location** | California, USA | San Diego, USA | California, USA | California, USA | South Carolina, USA |
| **Library prep kit** | Rapid barcoding kit 24 V14 (SQK-RBK114-24) | Rapid barcoding kit 24 V14 (SQK-RBK114-24) | Rapid barcoding kit 24 V14 (SQK-RBK114-24) | Rapid barcoding kit 24 V14 (SQK-RBK114-24) | Rapid barcoding kit 24 V14 (SQK-RBK114-24) |
| **Genome coverage (QC bp)** | 15.38 | 84.99 | 222.63 | 85.48 | 85.64 |
| **Total length (assembly bp)** | 45,852 | 45,254 | 44,997 | 46,094 | 50,563 |
| **CDS** | 86 | 66 | 63 | 64 | 84 |
| **Hypothetical proteins** | 45 | 39 | 27 | 37 | 48 |
| **VC taxonomy** | Heunggongvirae,Uroviricota,Caudoviricetes,Caudovirales,Siphoviridae,Steinhofvirus | Heunggongvirae,Uroviricota,Caudoviricetes,Caudovirales,Siphoviridae,Steinhofvirus | Heunggongvirae,Uroviricota,Caudoviricetes,Caudovirales,Siphoviridae,Steinhofvirus | Heunggongvirae,Uroviricota,Caudoviricetes,Caudovirales,Siphoviridae,Steinhofvirus | Heunggongvirae,Uroviricota,Caudoviricetes,Caudovirales,Siphoviridae,Steinhofvirus |
| **Taxa description** | Achromobacter phage phiAxp-1 | Achromobacter phage vB_AxyS_19-32_Axy16 | Achromobacter phage vB_AxyS_19-32_Axy18 | Achromobacter phage vB_AxyS_19-32_Axy16 | Achromobacter phage AMA1 |
| **Lowest taxa classification** | Unclassified | Steinhofvirus | Unclassified | Steinhofvirus | Steinhofvirus |
| **Matched hashes** | 78 | 423 | 480 | 414 | 406 |

**Footnote:** All phages produced circular, clear plaques with no halo formation, though plaque size varied among isolates. All phages were already visualised using TEM and classified within the *Siphoviridae* family and exhibited isometric heads. Measured head dimensions and tail lengths.

**Supplementary Table S6:** Linear-mixed effect models results for the fixed and interaction effects

| Main effects (95% HDI) | | | |
| --- | --- | --- | --- |
| Parameter | β | Highest density interval (HDI) 2.5% | HDI 97.5% |
| Host species; xylosoxidans | 0.564 | -30.776 | 31.88 |
| Phage family: Steinhofvirus | 0.793 | -32.865 | 34.70 |
| Defence subtype: AbiU | 0.017 | -43.453 | 42.88 |
| Defence subtype: CBASS II | -0.193 | -43.453 | 42.88 |
| Defence subtype: CBASS III | 0.017 | -29.519 | 29.757 |
| Defence subtype: Hna | -0.896 | -43.614 | 40.299 |
| Defence subtype: RM Type II | -0.101 | -33.325 | 33.137 |
| Defence subtype: Shedu | -0.193 | -31.804 | 31.628 |
| AMR Class: Aminoglycoside | -0.88 | -33.410 | 31.648 |
| AMR Class: Chloramphenicol | -0.255 | -33.578 | 33.522 |
| AMR Class: Fosfomycin | -0.033 | -36.276 | 36.444 |
| AMR Class: Tetracycline | 0.967 | -46.559 | 48.138 |
| AMR phenotype (mean) | 0.967 | -46.559 | 48.139 |
| Tail genes: Cluster 1 | -1.084 | -29.258 | 26.961 |
| Tail genes: Cluster 2 | 1.158 | -7.063 | 9.400 |
| Tail genes: Cluster 3 | -2.125 | -7.744 | 3.734 |
| Tail genes: Cluster 4 | -1.870 | -10.814 | 7.077 |
| Tail genes: Cluster 5 | 2.884 | -25.510 | 30.653 |
| Tail genes: Cluster 6 | -6.174 | -16.675 | 4.843 |
| Tail genes: Cluster 7 | -1.764 | -10.964 | 7.520 |
| Tail genes: Cluster 8 | -1.146 | -35.109 | 32.255 |
| INTERACTION EFFECTS (defence × cluster) | | | |
| def_AbiU:cl_cluster_1 | 1.755 | -43.808 | 45.795 |
| def_AbiU:cl_cluster_2 | 0.142 | -26.555 | 26.504 |
| def_AbiU:cl_cluster_3 | -2.126 | -36.201 | 30.778 |
| def_AbiU:cl_cluster_4 | -1.607 | -48.964 | 44.733 |
| def_AbiU:cl_cluster_5 | -6.215 | -68.093 | 53.979 |
| def_AbiU:cl_cluster_6 | 4.185 | -61.130 | 69.609 |
| def_AbiU:cl_cluster_7 | 1.638 | -46.207 | 48.795 |
| def_AbiU:cl_cluster_8 | 1.211 | -53.794 | 56.190 |
| def_CBASS_II:cl_cluster_1 | 0.346 | -32.907 | 35.335 |
| def_CBASS_II:cl_cluster_2 | -0.688 | -27.013 | 25.735 |
| def_CBASS_II:cl_cluster_3 | 0.112 | -32.452 | 33.909 |
| def_CBASS_II:cl_cluster_4 | 0.094 | -46.250 | 46.691 |
| def_CBASS_II:cl_cluster_5 | -3.122 | -55.274 | 49.365 |
| def_CBASS_II:cl_cluster_6 | 3.829 | -60.710 | 68.054 |
| def_CBASS_II:cl_cluster_7 | -0.112 | -47.259 | 47.199 |
| def_CBASS_II:cl_cluster_8 | 1.346 | -43.669 | 47.599 |
| def_CBASS_III:cl_cluster_1 | 1.129 | -26.550 | 29.235 |
| def_CBASS_III:cl_cluster_2 | 1.820 | -3.629 | 7.264 |
| def_CBASS_III:cl_cluster_3 | -2.780 | -6.450 | 0.862 |
| def_CBASS_III:cl_cluster_4 | -3.092 | -8.880 | 2.785 |
| def_CBASS_III:cl_cluster_5 | -2.687 | -31.203 | 25.695 |
| def_CBASS_III:cl_cluster_6 | -0.088 | -6.875 | 6.483 |
| def_CBASS_III:cl_cluster_7 | 0.163 | -5.650 | 5.973 |
| def_CBASS_III:cl_cluster_8 | -1.589 | -29.946 | 26.580 |
| def_Hna:cl_cluster_1 | 1.008 | -39.682 | 41.771 |
| def_Hna:cl_cluster_2 | 0.853 | -6.531 | 7.961 |
| def_Hna:cl_cluster_3 | -4.008 | -8.786 | 0.845 |
| def_Hna:cl_cluster_4 | -1.611 | -9.337 | 6.117 |
| def_Hna:cl_cluster_5 | -5.452 | -46.715 | 36.345 |
| def_Hna:cl_cluster_6 | 2.586 | -6.301 | 11.744 |
| def_Hna:cl_cluster_7 | 4.397 | -3.185 | 12.273 |
| def_Hna:cl_cluster_8 | -2.599 | -43.684 | 39.074 |
| def_RM_Type_II:cl_cluster_1 | -0.213 | -34.571 | 33.616 |
| def_RM_Type_II:cl_cluster_2 | 1.406 | -24.610 | 28.000 |
| def_RM_Type_II:cl_cluster_3 | -0.121 | -32.961 | 33.891 |
| def_RM_Type_II:cl_cluster_4 | 0.726 | -45.894 | 48.149 |
| def_RM_Type_II:cl_cluster_5 | -1.123 | -53.352 | 51.869 |
| def_RM_Type_II:cl_cluster_6 | 0.450 | -64.075 | 64.847 |
| def_RM_Type_II:cl_cluster_7 | -0.317 | -46.708 | 47.121 |
| def_RM_Type_II:cl_cluster_8 | 0.225 | -46.340 | 46.186 |
| def_Shedu:cl_cluster_1 | -0.610 | -31.186 | 30.450 |
| def_Shedu:cl_cluster_2 | 0.609 | -25.654 | 26.646 |
| def_Shedu:cl_cluster_3 | 0.164 | -33.411 | 32.814 |
| def_Shedu:cl_cluster_4 | 1.179 | -45.152 | 47.776 |
| def_Shedu:cl_cluster_5 | 0.062 | -50.355 | 50.967 |
| def_Shedu:cl_cluster_6 | 1.061 | -63.290 | 65.996 |
| def_Shedu:cl_cluster_7 | 0.309 | -47.695 | 46.773 |
| def_Shedu:cl_cluster_8 | 0.433 | -43.994 | 45.091 |

### Supplementary Figures

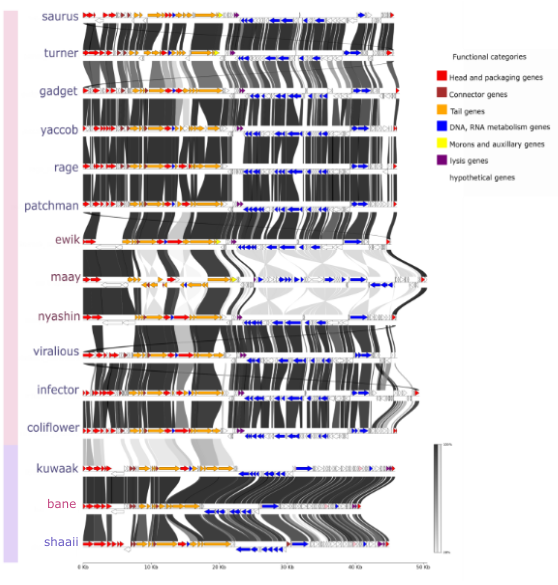

**Supplementary Figure S1:** Gene-by-gene comparison across the 15 *Achromobacter* phages, with genes colour coded by PHROG functional category. Connecting lines denote pairwise gene similarities ranging from 0% (white) to 100% (black). The bars to the right represent pink for taxa cluster 1 and purple representing genomes for taxa cluster 2.

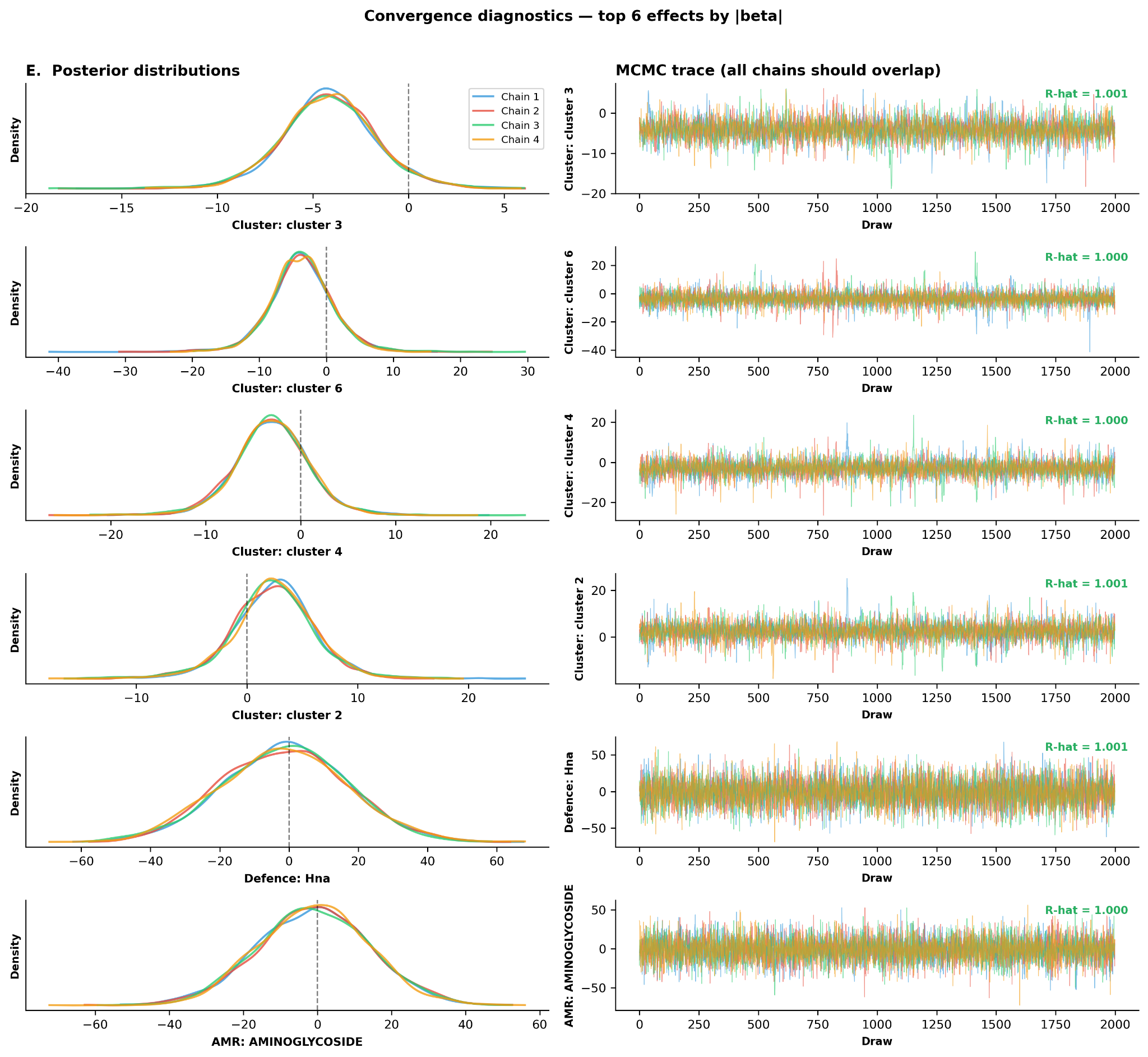

##### **Supplementary Figure S2: Convergence diagnostics for the Bayesian linear mixed-effects model.**Trace plots and posterior distributions are shown for the six fixed-effect parameters with the largest absolute posterior mean (β) from the Bayesian linear mixed-effects model. For each parameter, the left panel shows the posterior density estimated by kernel density estimation (KDE) separately for each of the four independent MCMC chains (Chain 1–4, colour-coded), and the right panel shows the corresponding MCMC trace — the sampled parameter values across all 2,000 posterior draws per chain. The vertical dashed line in the left panels indicates zero. The R-hat statistic (Gelman–Rubin convergence diagnostic) is annotated in the upper right of each trace panel; values below 1.01 indicate adequate convergence of the four chains to a common posterior distribution. All six parameters shown have R-hat ≤ 1.002, confirming that the sampler explored the posterior reliably and that parameter estimates are not artefacts of poor mixing. The wide posterior distributions and large credible intervals visible in both panels are a consequence of the limited sample size (n = 105, seven bacterial strains, fifteen phages) relative to model complexity, and reflect genuine uncertainty in the data rather than sampling failure. The complete model included host species, phage family, six defence system subtypes, four AMR gene classes, mean AMR phenotype score, eight phage tail clusters, and 48 defence × cluster interaction terms as fixed effects, with crossed random intercepts for bacterial strain (1 | bacteria) and phage identity (1 | phage_id). The model was fitted using the No-U-Turn Sampler (NUTS) implemented in PyMC (v5) via the bambi interface (v0.14), with 4 chains, 2,000 tuning steps, and 2,000 posterior draws per chain (8,000 total samples), a target acceptance rate of 0.99, and a maximum tree depth of 12.
